# Nonadditive gene expression and reduced homoeolog expression bias in an intraspecific hexaploid wheat hybrid

**DOI:** 10.64898/2026.06.29.735383

**Authors:** Asena Ardaman, Cristiane Forgiarini, Ramesh Arunkumar

**Affiliations:** Section of Population Genetics. Hans-Eisenmann-Zentrum für Agrarwissenschaften. School of Life Sciences. Technical University of Munich. Liesel-Beckmann-Strasse 2. 85354 Freising, Germany; Istanbul University-Cerrahpaşa, Kocamustafapaşa Cad. No:53, 34098 Fatih, Istanbul, Turkey; UK Centre for Ecology and Hydrology, Wallingford, Oxfordshire, OX10 8BB, United Kingdom

**Keywords:** Triticum aestivum, allopolyploidy, transcriptome, bisulfite sequencing, gene body methylation, transgressive expression, triads, F₁ hybrids

## Abstract

Intraspecific hybridization in allopolyploid plants can generate additive and nonadditive changes in gene expression through interactions between divergent parental genomes. However, how it simultaneously affects gene expression and the relative expression of homoeologs in higher-order polyploids is less well understood. To study this, we sequenced seedling leaf transcriptomes and profiled gene body methylation in two hexaploid wheat (*Triticum aestivum* L.) cultivars and their F₁ hybrids. Although only 4.3% of genes differed in expression between the parents, 22.3% deviated from mid-parent expression in the hybrids, with many showing transgressive expression. 32.1% of triads contained at least one homoeolog that deviated from mid-parent expression, and all three homoeologs deviated in 11% of triads, substantially more than expected by chance. Triads in which all three homoeologs were overexpressed also showed reduced differences in expression among homoeologs. Greater parental divergence in relative homoeolog expression was associated with nonadditive expression. Genes lacking gene body methylation were also more likely to show dominant or transgressive expression, whereas gene body methylation was associated with more balanced homoeolog expression and additive or conserved expression. Intraspecific hybridization in hexaploid wheat, even without a change in ploidy, was associated with widespread nonadditive gene expression and altered relative homoeolog expression within triads. These responses were associated with parental differences in homoeolog expression and the absence of gene body methylation.

## Background

Hybridization between cultivars is an important strategy in crop breeding for developing varieties with beneficial traits[1], but hybrid performance depends on how parental genomes interact. In newly formed hybrids, interactions among locally acting *cis* genetic variants, distantly acting *trans* genetic variants, epigenetic marks, and/or transposable elements can lead to “genomic shock”, in which many genes are differentially expressed in hybrid offspring relative to their parents[2–4]. Polyploidy introduces another layer of complexity when studying the impact of hybridization on gene expression. The presence of paired genes that diverged after speciation and were later brought back together in the same genome through allopolyploidization, known as homoeologs[5], can provide evolutionary flexibility and robustness. Some homoeologs may acquire new functions, become subfunctionalized, or buffer phenotypic effects by acting redundantly[6]. This feature has been particularly important in the domestication and improvement of many crop species[7]. Hybridization and allopolyploidization create opportunities for novel *cis*–*trans* regulatory interactions, as parental alleles and homoeologs are exposed to a shared pool of *trans*-acting regulators within the same nucleus[8], In allotetraploid *Brassica napus* and cotton, intraspecific hybridization is associated with additive and nonadditive changes in gene expression and reduced expression divergence between homoeologs[9–12], but whether this pattern extends to higher-order polyploids remains unclear. Hexaploid wheat provides an opportunity to test how interactions among three subgenomes shape gene expression and relative homoeolog expression following hybridization.

*Triticum aestivum* L. (2n = 6x = 42; BBAADD) is a domesticated allohexaploid crop that originated through two hybridization and polyploidization events. The first occurred approximately 0.5–0.8 million years ago, when the A-genome donor *Triticum urartu* hybridized with an unknown B-genome donor, giving rise to tetraploid wild emmer wheat[13]. The second occurred approximately 8,000–11,000 years ago, when domesticated tetraploid wheat hybridized with the D-genome donor *Aegilops tauschii*, giving rise to hexaploid bread wheat[13]. As a result, many genes in bread wheat are present as three related homoeologous copies, one on each of the A, B, and D subgenomes[14]. A set of three homoeologs is commonly referred to as a triad[14]. Although many triads retain balanced expression among the three subgenomes, about 30% show homoeolog expression bias (HEB), where one homoeolog contributes disproportionately to the total triad expression[14].

Intraspecific hybridization generates additive and nonadditive gene expression patterns in hexaploid wheat. In Jingmai 8 seedlings and spikes, nonadditive genes, defined as those whose expression in the hybrid differed from the mid-parental value, accounted for 1.0–1.6% of expressed genes and 18.7–33.4% of genes differentially expressed between the hybrid and at least one parent[15]. Most of these changes reflected similarity to one parent or the other[15], a pattern termed parental expression-level dominance. A more recent study of the heterotic hybrid BC98 found that more than 50% of genes showed nonadditive expression in flag leaves at anthesis and during grain filling, although the extent of nonadditivity decreased during development[16]. Again, nonadditive expression primarily reflected parental expression-level dominance. The same study found that *cis*-only regulatory effects were more prevalent at the later developmental stage, whereas *trans*-regulatory effects contributed more strongly at the earlier stage[16]. For selected yield-related triads, Awan et al.[17] found that HEB varied among parental hexaploid wheat cultivars and their F₁ hybrids. Although this study demonstrates that HEB depends on genetic background[17], it was limited to a subset of genes. An unresolved question is therefore how parental gene expression differences and expression changes following intraspecific hybridization influence HEB in hexaploid wheat.

Another important question concerns the mechanisms generating these expression changes. Both *cis*- and *trans*-regulatory variation contribute to HEB in bread wheat[18–21], suggesting that gene regulatory divergence between parents may contribute to HEB patterns in their hybrids. In addition, parental HEB differences may also reflect homoeolog-specific expression divergence that predisposes genes to altered homoeolog expression in F₁ hybrids. Another potential influence is DNA cytosine methylation, an epigenetic mark frequently associated with gene regulation. However, cytosine methylation at promoters and transcription start sites shows weak or no association with gene expression or HEB in hexaploid wheat[22, 23]. In contrast, gene body methylation (gbM), a conserved epigenetic state characterized by high CG methylation and low CHG and CHH methylation within gene bodies[24], is associated with stable expression, broad expression across tissues, and balanced homoeolog expression[14]. CG methylation is also transgenerationally stable, making it useful for investigating the evolutionary potential of plant populations[25–27]. Regulatory divergence between parents, parental HEB differences, and gbM status may therefore influence changes in gene expression and HEB following intraspecific hybridization.

We profiled seedling leaf transcriptomes and methylomes from Chinese Spring (CS), Paragon, and their F_1_ hybrid. These cultivars are important genetic resources for bread wheat: CS is the primary bread wheat reference genotype[28], whereas Paragon is a spring elite cultivar extensively used for association mapping[29]. We investigated whether intraspecific hybridization is associated with simultaneous expression shifts among homoeologs within triads and whether parental regulatory divergence, parental HEB differences, and gbM are associated with changes in gene expression and HEB in hybrids relative to their parents.

## Results

### Transgressive gene expression in intraspecific hexaploid wheat hybrids

We first compared gene expression between parental *T. aestivum* cultivars and their hybrids. We sequenced transcriptomes from three biological replicates of the Paragon and CS parents and their reciprocal hybrids. Principal component analysis (PCA) of gene expression counts showed genotype-specific clustering, with P×CS3 as the only outlier (Fig. S1), suggesting a sample-specific transcriptomic effect. We therefore excluded this sample from subsequent analyses. Only two genes differed significantly in expression between the reciprocal crosses (P×CS vs CS×P; false discovery rate (FDR)<0.05; Fig. S2). We excluded these genes and combined the remaining reciprocal-cross samples into a single hybrid group. Repeating the PCA showed clear genotype-level clustering along the first two principal components (Fig. 1A).

**Figure 1.**
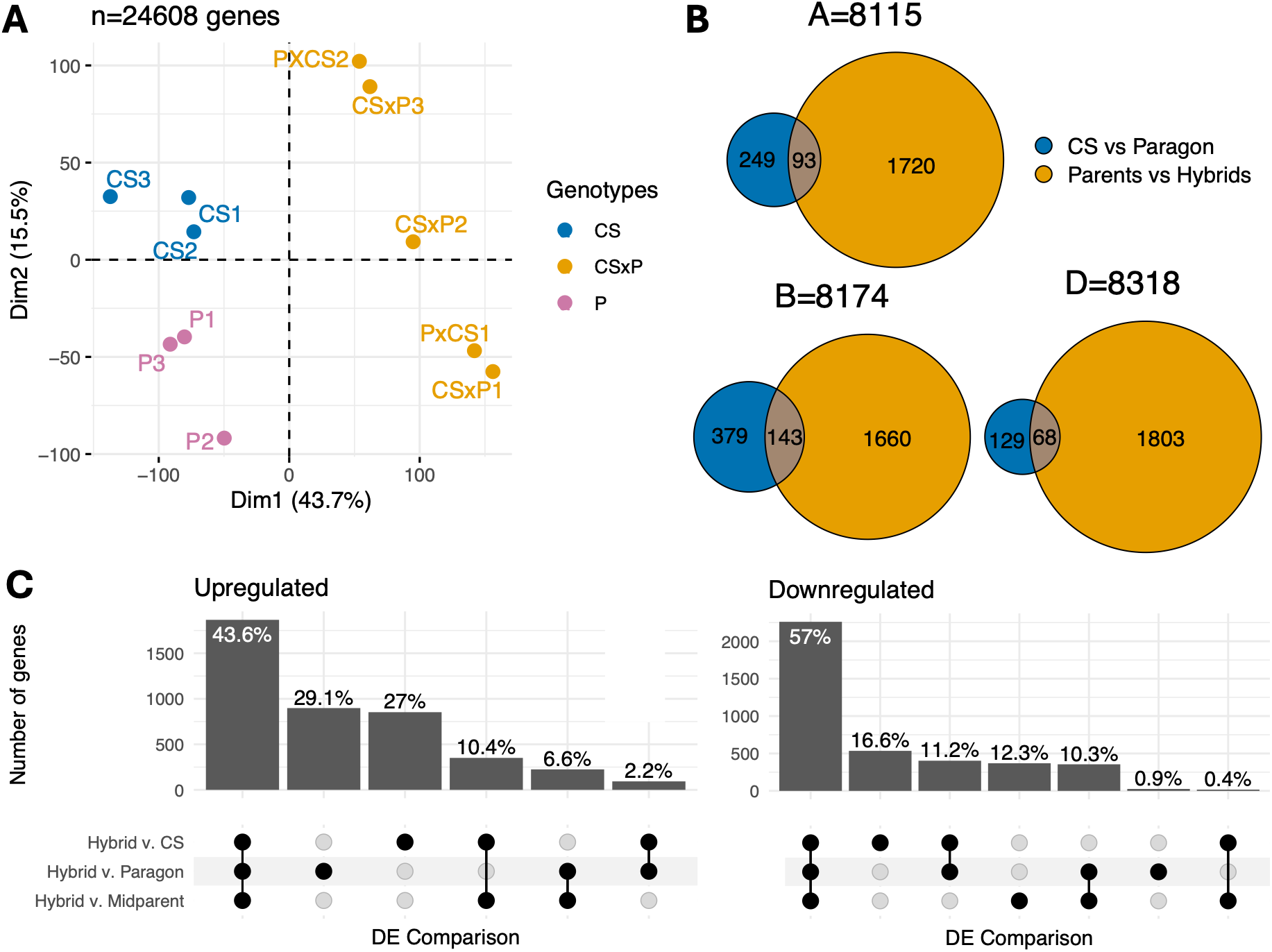
Nonadditive gene expression following intraspecific hybridization in *Triticum aestivum*. **(A)** Principal component analysis based on read counts for Chinese Spring (CS), Paragon (P), and hybrid genotypes. **(B)** Number of differentially expressed genes between CS and Paragon, and between hybrids and mid-parent estimates. The number of detected or classified genes in each subgenome is shown. **(C)** Intersections of differentially expressed (DE) genes upregulated or downregulated in hybrids relative to CS, Paragon, and the mid-parent expectation. For (B) and (C), DE was identified using a limma–voom empirical Bayes moderated t-test (≥2-fold expression difference and FDR<0.05).

Next, we compared gene expression patterns in the parents and hybrids. We found that 4.3% of the 24,608 detected genes were differentially expressed between CS and Paragon, whereas 22.3% differed in expression between the hybrids and mid-parent expectations, using thresholds of ≥2-fold expression difference and FDR<0.05 (Fig. 1B; Fig. S3). The proportion of differentially expressed genes between the parents differed significantly among subgenomes (A=4.2%, B=6.4%, D=2.4%; χ²=161.2, df=2, *P*<0.001), whereas the proportion of genes differentially expressed between hybrids and mid-parent expectations did not (χ²=0.46, df=2, *P*=0.79), indicating that hybridization affected all subgenomes similarly. Given the large number of genes deviating from mid-parent expectations, we tested whether hybrid expression primarily reflected dominance toward one parent. Instead, 43.6% of the genes upregulated in the hybrids relative to the mid-parent value were also upregulated when the hybrid was compared with the individual parental lines (Fig. 1C). Similarly, 57% of the genes downregulated in the hybrids relative to the mid-parent value were also downregulated when the hybrid was compared with the individual parental lines (Fig. 1C), indicating that hybridization shifted expression beyond the parental values.

We tested for allele-specific expression (ASE) in the hybrids. Because genomic DNA sequencing data were unavailable to estimate allelic ratios while accounting for mapping bias, we used a conservative approach (detailed in Methods). Briefly, we retained only 1:1 orthologs, defined here as the corresponding genes identified from separate cultivar assemblies, between CS and Paragon for which parental RNA-seq reads mapped uniquely and without mismatches to the corresponding haplotype in a combined transcriptome reference. Among the 770 retained genes, we found no evidence of genome-wide mapping bias toward either parental haplotype (Fig. S4). We excluded 12 genes showing reciprocal-cross-dependent ASE (Fig. S5), and 6.5% of the remaining genes showed significant ASE (≥2-fold difference, FDR < 0.05; Fig. S6). We then classified regulatory divergence using the widely used approach of McManus *et al*.[30]. Under this approach, 62.5% of 717 genes were classified as conserved, 27.1% ambiguous, 1% *cis*-only, and 5.7% *trans*-only (Fig. S7A). Because this approach may overestimate *trans* effects when expression levels varies among individuals, we also applied a linear-model approach adapted from Haas *et al*.[31]. This classified 83.1% of genes as conserved, 13.5% as ambiguous, 2.6% as *cis*-only, and 0.1% as *trans*-only (Fig. S7B). The limited parental expression divergence, stringent filtering, and individual-level variation prevented us from identifying clear patterns of parental regulatory divergence.

### Hybridization associated with triad-wide expression changes

As differential expression among wheat homoeologs derived from separate progenitor species may help regulate protein dosage[14], we tested how hybridization affected homoeolog expression within triads. We identified 6,591 triads where all three homoeologs were detected in the expression dataset. Among these, 10.7% had at least one homoeolog differentially expressed between CS and Paragon, usually involving only one of the three homoeologs (Fig. 2A). In contrast, 32.1% of triads had at least one differentially expressed homoeolog between the hybrids and the mid-parent expression values. Interestingly, all three homoeologs were differentially expressed in 727 triads (Fig. 2B). Because hybridization was associated with widespread differential expression (Fig. 1B), we tested whether this pattern could arise by chance. Only ∼73 triads were expected to contain three differentially expressed homoeologs by chance alone (Fig. S8), suggesting that the observed pattern is unlikely to be purely random. We then measured HEB in the parents and hybrids as the coefficient of variation of the relative expression proportions of homoeologs within each triad, as described previously[19]. For triads in which all three homoeologs were overexpressed relative to the parents (Fig. S9A), hybrid HEB was lower than in both parents (Fig. 2C), suggesting that hybridization results in more balanced homoeolog expression. This trend was not observed for triads in which all three homoeologs were underexpressed in hybrids (Fig. S9B–C).

**Figure 2.**
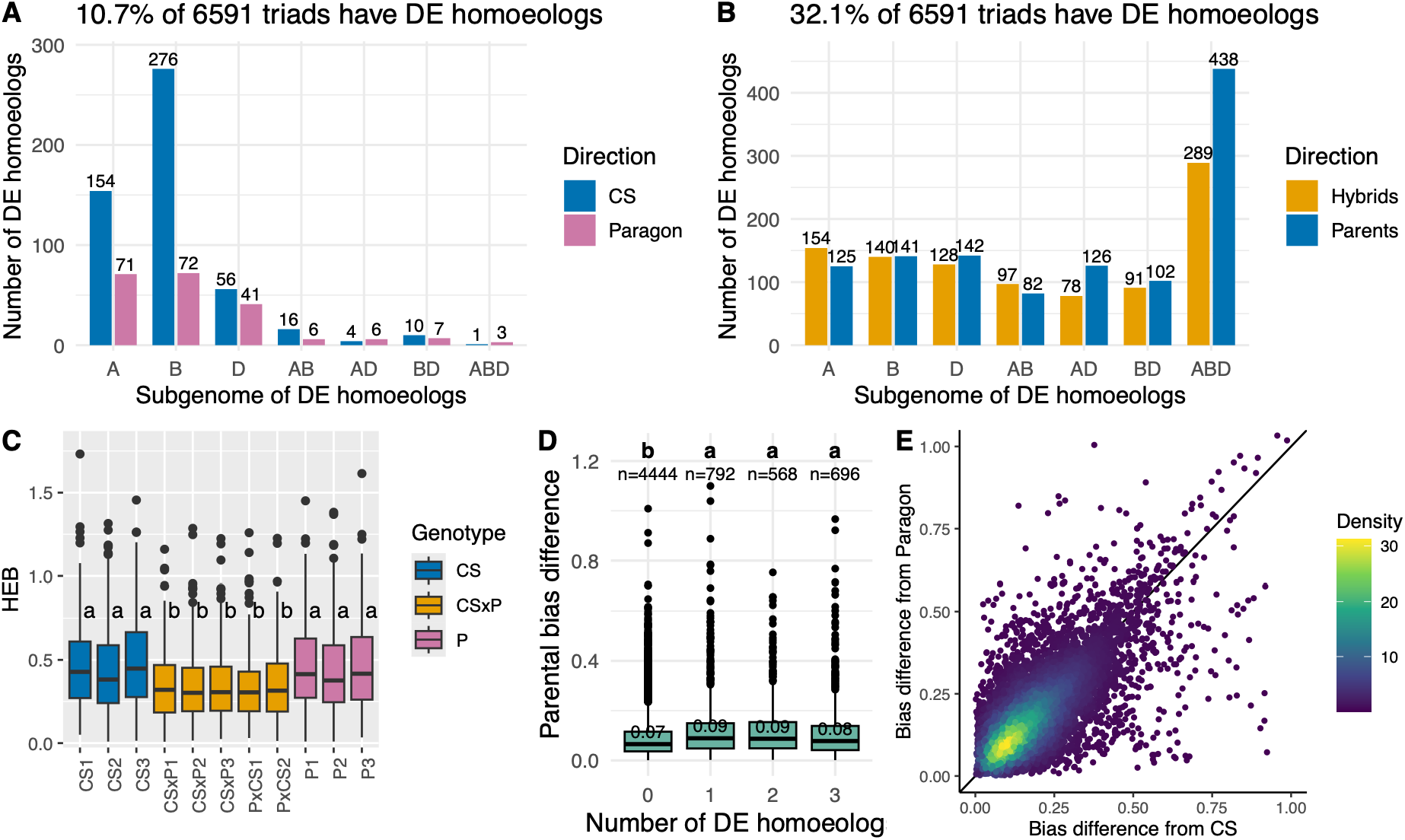
Parental differences in homoeolog expression bias (HEB) are associated with differential homoeolog expression following intraspecific hybridization in *Triticum aestivum*. **(A)** Frequency of homoeologs differentially expressed (DE) between Chinese Spring (CS) and Paragon. **(B)** Frequency of homoeologs DE between hybrids and mid-parent expression. For (A-B), direction indicates the sample in which expression was higher; the numbers are the counts of triads in each category; DE was identified using a limma–voom empirical Bayes moderated t-test (≥2-fold expression difference and FDR<0.05). **(C)** Homoeolog expression bias (HEB), measured using the coefficient of variation of the relative homoeolog expression proportions, for triads where all three homoeologs are overexpressed in hybrids compared to the parents. Numbers are median values. **(D)** Association between parental HEB divergence, measured using Euclidean distances, and the number of homoeologs in a given triad that are DE in hybrids. Numbers in boxes are median values. n refers to the number of triads. For (C-D), letters indicate Tukey’s honestly significant difference groupings following one-way analysis of variance; categories sharing a letter do not differ significantly (*P*>0.05). **(E)** Hybrid HEB divergence from CS and Paragon for triads with homoeologs DE in hybrids compared to parents. Points represent triad values pooled across the five hybrid genotypes.

We next investigated whether parental HEB differences were associated with expression changes and HEB in hybrids. We measured HEB divergence between CS and Paragon as the Euclidean distance between their relative homoeolog expression proportions, which sum to 1, following the approach in[19]. Triads with greater parental HEB divergence were more likely to contain at least one differentially expressed homoeolog in hybrids (Fig. 2D). When gene expression changed in hybrids compared to the mid-parent expression, hybrid HEB predominantly deviated from both parents on a linear scale (Fig. 2E), indicating distinct bias patterns in hybrids. Parental HEB divergence was associated with whether HEB changed following hybridization, but not with the direction of that change.

### Gene body methylation associated with additive expression and lower homoeolog expression bias

We next asked whether differential gene expression in the hybrid was associated with the presence or absence of gbM. Using sites covered by at least 10 reads (Fig. S10), our estimates of mean genome-wide methylation ranged from 81–84% at CG sites, 53–61% at CHG sites, and 2–3% at CHH sites (Fig. S11), consistent with previous estimates from wheat leaves[32, 33]. Whereas genome-wide methylation in *T. aestivum* is strongly influenced by the abundance of transposable elements[34], mean coding sequence methylation was lower, at 19–20% for CG, 2–3% for CHG, and 1–2% for CHH sites (Fig. S11). We then inferred gbM status by testing whether genes were enriched for methylated CG sites, but not CHG or CHH methylation, relative to genotype- and context-specific coding sequence (CDS)-wide background frequencies. Among classifiable genes, 39% were gbM. The frequency of gbM genes did not differ significantly among the A, B, and D subgenomes (χ²=2.7, df=2, *P*=0.26). gbM was not associated with gene expression level (Fig. 3A) but was associated with lower HEB (Fig. 3B). We then compared gbM frequencies among additive (hybrid expression matched the mid-parent value), dominant (hybrid expression matched one parent), transgressive (hybrid expression fell outside parental range), and non-DE (no significant expression differences among parents and hybrids) genes. Genes with nonadditive expression in the hybrid, classified as transgressive or dominant, were less likely to have gbM than additive or non-DE genes (χ²=199, df=3, *P*<0.001; Fig. 3C). gbM was associated with more conserved and additive gene expression and balanced homoeolog expression.

**Figure 3.**
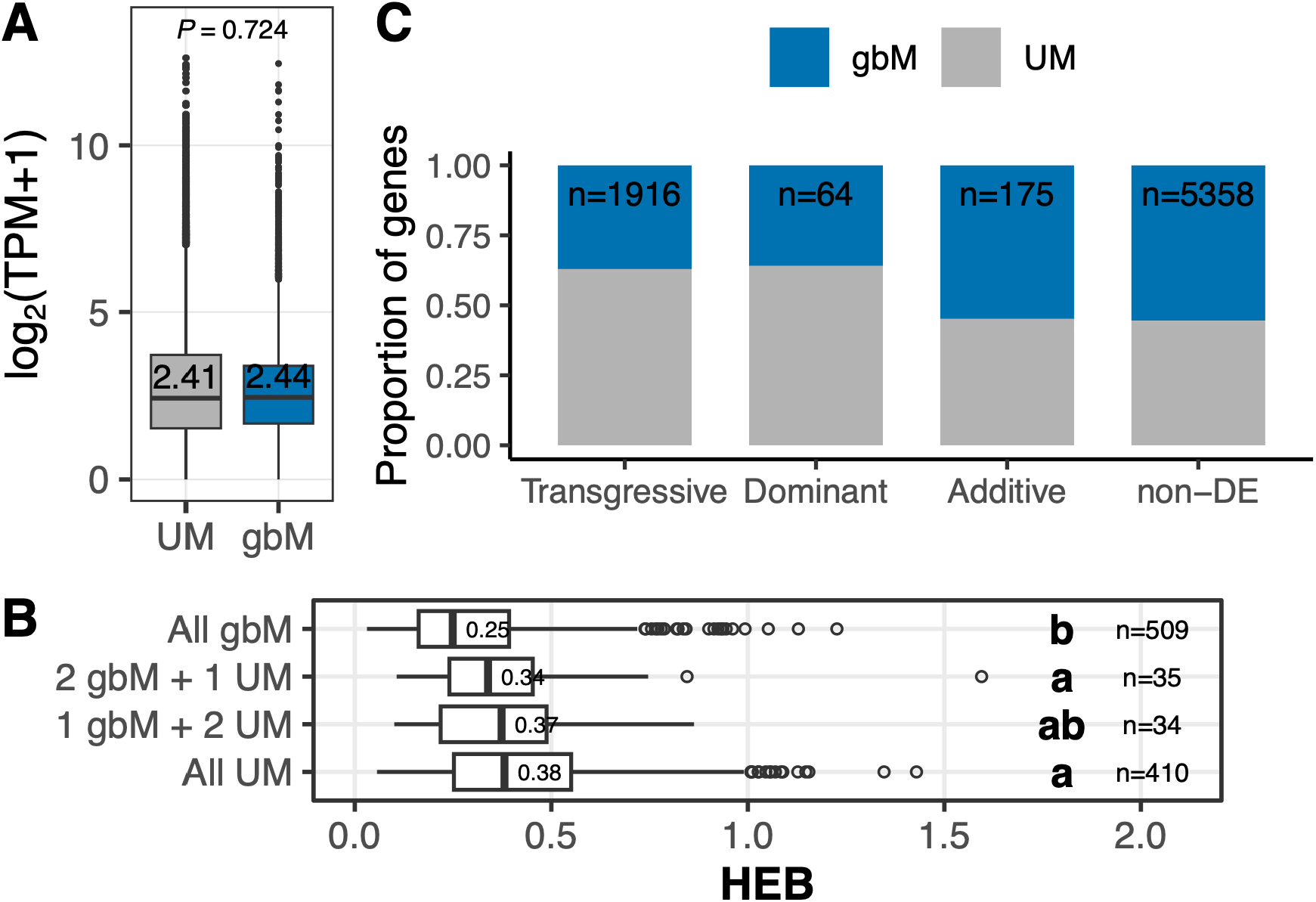
Gene body methylation (gbM) associated with differential gene and homoeolog expression following intraspecific hybridization in *Triticum aestivum*. **(A)** Expression levels of gbM and unmethylated (UM) genes, measured as transcripts per million (TPM). Wilcoxon test *P*-values and medians are shown. **(B)** Homoeolog expression bias (HEB), measured using the coefficient of variation of the relative homoeolog expression proportions, in triads with different numbers of gbM genes. Letters indicate significant differences based on Tukey’s honestly significant difference test following one-way analysis of variance; categories sharing a letter are not significantly different (*P*>0.05). Median values are shown. n indicates the number of triads per category. For (A-B), expression estimates from the parents and hybrids were averaged by genotype. **(C)** Proportion of gbM among transgressive, dominant, additive, and non-DE genes. Non-DE genes did not differ in parent–parent or parent–hybrid comparisons; additive genes matched the mid-parent expression; dominant genes matched the expression of one parent; transgressive genes were expressed above or below both parents. n indicates the number of genes per category. In all cases, only genes with identical gbM states in both parents and the hybrid were included.

## Discussion

We show that intraspecific hybridization in hexaploid bread wheat can reshape gene expression and HEB. Although only 4.3% of detected genes differed in expression between the parents, 22.3% differed between the hybrid and the mid-parent value, indicating widespread nonadditive expression following hybridization. These changes were frequently transgressive and often involved entire triads. We also tested whether differential gene expression in the hybrid was associated with three factors. The first was parental regulatory divergence. Although 6.5% of genes (50 genes) exhibited ASE, the limited expression divergence between the parents, together with our stringent filtering that retained only one-to-one orthologs, resulted in relatively few informative genes and limited our ability to quantify the contribution of regulatory divergence. The second factor was parental HEB. Triads with greater parental differences in HEB were more likely to contain at least one homoeolog that was differentially expressed in the hybrid. We previously showed that small parental HEB differences (<0.2) increase the likelihood that offspring HEB diverges similarly from both parents[19], which may explain the comparable hybrid–parent distances observed for HEB in this study. The third factor was gbM, which was associated with more balanced homoeolog expression but not with overall gene expression levels. This pattern is consistent with a role for gbM in stabilizing gene expression levels[14]. Genes lacking gbM were more likely to exhibit nonadditive expression in the hybrid. Our results suggest that parental HEB and gbM influence how genes respond transcriptionally to intraspecific hybridization in hexaploid wheat.

### Widespread nonadditive transcriptomic changes in an intraspecific wheat hybrid

We detected levels of nonadditive expression in wheat hybrids that were broadly comparable to a previous flag-leaf study of winter wheat hybrids at anthesis and grain filling[16]. In that study, nonadditively expressed genes were dominated by parental expression-level dominance[16], whereas we detected a much larger contribution of transgressive changes. This difference may partly reflect lower parental expression divergence in our study, because a narrower parental expression range can make hybrid shifts more likely to fall outside the parental range. Developmental stage may also contribute, as the previous study reported that additive, *cis*, and compensatory effects increased during development[16]. A second winter wheat study detected much lower parent-parent and hybrid-parent expression divergence, and nonadditive genes represented only a small fraction of expressed genes in seedlings[15]. In that cross, one parent is a thermosensitive male-sterile line whose sterility is associated with altered expression and methylation of reproductive and metabolic pathways[35]. This comparison further suggests that specific parental genotypes and developmental context influence how expression is remodeled after intraspecific hybridization in wheat.

Triads in which all three homoeologs were nonadditive and overexpressed in hybrids had lower HEB, raising the possibility that their overexpression has a shared regulatory basis. Similar patterns have been reported in other hybrid systems: homoeologs often changed expression in the same direction in *Arabidopsis* allotetraploids[36], expression divergence among homoeologs was reduced in *Brassica* and cotton hybrids relative to their parents[9, 37], and a standardized cross-species comparison found that genes differing in expression between parents often became more similarly expressed in plant, animal, and fungal hybrids[38]. One proposed mechanism is *trans*-regulatory cross-talk, whereby factors encoded by the two parental genomes can act on alleles derived from both parents within the shared hybrid nucleus[36, 38]. Although we cannot demonstrate coordinated regulation in bread wheat, it is plausible because many wheat triads show similar or correlated expression across tissues and conditions[14, 39], and therefore warrants further investigation.

Nonadditive genes had lower gbM, suggesting that genes lacking gbM may be more responsive to the altered regulatory environment created by hybridization. Canonical, stable gbM is associated with housekeeping genes and reduced expression variability across cells and conditions, while a partial loss of gbM in *Arabidopsis METHYLTRANSFERASE1* (*MET1-1*) mutant increased transcriptional noise[24, 40]. In bread wheat, progressive loss of *MET1-1* homoeologs caused dosage-dependent reductions in CG methylation and stepwise increases in the number of differentially expressed genes[32]. Together, these findings raise the possibility that genes with lower gbM are less transcriptionally buffered against changes in the regulatory environment of the hybrid genome. Future studies using targeted epigenetic manipulation could directly test this hypothesis.

### Limitations of our study

Our study characterizes changes in gene expression and HEB in an intraspecific wheat hybrid, but the observed patterns may not be generalizable to other wheat crosses. Wheat cultivars differ substantially in gene content, with approximately one-third of genes showing presence– absence variation among nine wheat cultivars[41]. Our findings are also limited to seedling leaves. Testing additional tissues and developmental stages could determine whether these patterns are associated with tissue-specific heterotic traits. Our conservative ASE analysis included only one-to-one orthologs between Paragon and CS and retained only uniquely mapped reads to minimize mapping bias. However, this excluded duplicated and highly similar genes. Because duplicated genes can undergo regulatory and functional divergence and are often associated with tissue-specific or stress-responsive functions[42], they are important candidates for regulatory divergence that our analysis could not assess. Finally, our study identified associations rather than causal relationships. Despite these limitations, our results provide a basis for testing whether nonadditive expression and reduced HEB following intraspecific hybridization are generalizable across wheat cultivars, tissues, and developmental stages, and for experimentally validating these patterns.

## Conclusions

We identified widespread nonadditive gene expression in an intraspecific hexaploid wheat hybrid. These changes were associated with differences in HEB between the parents and with gbM status. Triads where all three homoeologs were overexpressed also showed reduced HEB, indicating that changes in expression can coincide with shifts in relative homoeolog contributions. Although too few informative genes were available to quantify *cis*- and *trans*-regulatory effects, both are known to influence hybrid expression and HEB[16, 18–21]. Other mechanisms not examined here, including transposable element variation, promoter methylation, chromatin state, and small RNAs, have also been implicated in hybrid misexpression[2] and warrant further study in intraspecific bread wheat hybrids. Profiling these mechanisms and nonadditive expression across tissues, developmental stages, and hybrid combinations alongside heterosis phenotypes will help reveal which regulatory changes are associated with hybrid performance in bread wheat.

## Materials and Methods

### Growth and Crossing of Parental Lines

We vernalized Paragon and CS seeds for 30 days before transferring them to the John Innes Centre (JIC) glasshouses. We germinated seeds on moist filter paper at 4°C for two days, followed by two days at room temperature, and then transferred the seedlings to pots containing JIC Cereal Compost Mix[43]. After spike emergence, we performed reciprocal crosses in April 2022 using the detached spike pollination method (https://www.wheat-training.com/). We emasculated pollen-recipient spikes before anther maturation and removed the apical and basal spikelets, which developed asynchronously relative to the central spikelets. We trimmed donor spikes to expose anthers and gently warmed them by hand to promote dehiscence. We then inverted donor spikes and enclosed them with recipient spikes in plastic bags. We gently tapped donor spikes to transfer pollen, allowed spikes to mature, harvested grains, and collected seeds after threshing.

### Seedling Growth, Nucleic Acid Extraction, and Sequencing

We germinated parental and hybrid seedlings as described above, but omitting vernalization. We grew seedlings in seed trays in the JIC glasshouses in February 2025 under 20°C day/16°C night conditions and a 16:10 h light:dark photoperiod. We potted seedlings in a 9:1 mixture of Levington F2 Starter + Grit compost and horticultural grit to improve drainage, with no additional fertilizer. Three weeks after germination, we collected leaf tissue from each seedling and flash-froze samples in liquid nitrogen. We extracted RNA using the Qiagen RNeasy Plant Mini Kit and DNA using the Qiagen DNeasy Plant Mini Kit, then stored nucleic acid extracts at −80°C before sequencing. We prepared poly(A)-enriched messenger RNA libraries and sequenced them with 150bp paired-end reads on an Illumina NovaSeq X Plus at Novogene GmbH, Munich, Germany, targeting ∼10Gb of raw data per sample. For methylation sequencing, equal quantities of DNA from three biological replicates were combined, yielding a single pooled biological replicate for each genotype. Three independent whole genome bisulfite sequencing libraries were prepared from each pooled replicate and sequenced using 150-bp paired-end reads on the DNBSEQ platform at Sailgene Technology, Hong Kong, targeting approximately 430Gb per genotype. These libraries were technical replicates derived from the same pooled biological sample, and their reads were combined for downstream analysis. We used the same leaf tissue as for RNA-seq but sequenced only hybrids from the CS×P cross.

### Mapping and Counting RNA-seq Reads

We followed the methods established in Glombik *et al*[19]. We processed raw reads using Trimmomatic version 0.39[44]. We removed adapter sequences (ILLUMINACLIP: two seed mismatches; palindrome and simple clip thresholds of 30 and 10, respectively), trimmed low-quality bases (quality score <3) from read ends, and applied a 4-bp sliding window to remove regions with average quality scores below 20. We discarded reads shorter than 36bp and unpaired reads. We quantified transcript abundance with Kallisto version 0.51.1[45] by mapping to the International Wheat Genome Sequencing Consortium (IWGSC) RefSeq v2.1[46] and summarized transcript-level estimates to gene-level counts with tximport version 1.36[47].

### Differential Gene Expression

We restricted all gene-centered analyses to core genes, which are shared among divergent wheat cultivars[41] and therefore facilitate cross-cultivar comparisons. We performed differential expression analysis with edgeR version 4.6.3[48], using high-confidence genes with counts per million (CPM) ≥2 in at least four samples. We calculated library sizes from the filtered counts and estimated normalization factors using the trimmed mean of M-values (TMM) method. We fitted a no-intercept design matrix to model group effects, applied the voom transformation[49], and performed linear modelling with empirical Bayes moderation using limma version 3.68.4[50]. We defined parental expression divergence from the CS– Paragon contrast using absolute log₂ fold change (|logFC|) >1 and FDR-adjusted *P*<0.05. For hybrid–mid-parent comparisons, we generated arithmetic mid-parent pseudo-replicates by averaging TMM-normalized counts from paired CS and Paragon replicates, then applied voom and fitted a separate hybrid–mid-parent model following the approach described in[51]. We visualized selected CPM values using pheatmap version 1.0.13[52] and ComplexHeatmap version 2.24.1[53], plotted overlaps between differential expression contrasts using eulerr version 7.0.4[54], and intersections between differentially expressed genes using the UpSet method[55].

Transgressive genes were defined as those with |logFC| >1 and FDR-adjusted *P*<0.05 in the hybrid–mid-parent comparison and in both hybrid–parent comparisons, with the hybrid showing either higher expression than both parents or lower expression than both parents. We classified genes as dominant when the parents differed in expression (|log₂FC| >1 and FDR-adjusted *P*<0.05), the hybrid differed from both the arithmetic mid-parent and one parent (|log₂FC| >1 and FDR-adjusted *P*<0.05), but not from the other parent (FDR-adjusted *P*≥0.05). The hybrid–mid-parent difference was also required to be in the direction of the parent that the hybrid resembled. We classified genes as additive when the parents differed in expression (|log₂FC| >1 and FDR-adjusted *P*<0.05), but the hybrid did not differ from the arithmetic mid-parent (FDR-adjusted *P*≥0.05). Non-DE genes were defined as those with FDR-adjusted *P*≥0.05 in the CS–Paragon, hybrid–mid-parent, and both hybrid–parent comparisons.

### Identification of One-to-One Orthologs

We used OrthoFinder version 2.5.5[56] to identify one-to-one orthologs between transcript annotations for high-confidence genes from the IWGSC RefSeq v2.1 CS reference and transcripts from the GCA949126075v1 Paragon reference from Ensembl Plants[57]. We analyzed the A, B, and D subgenomes separately. We ran BLASTN searches with an E-value threshold of 0.001, DendroBLAST gene-tree inference, and the default MCL inflation parameter of 1.5.

### Allelic Expression Quantification

We grouped transcript records from the IWGSC RefSeq v2.1 messenger RNA and Paragon GCA949126075v1 reference transcriptomes by gene and retained the longest transcript sequence per gene. We concatenated these transcripts into a combined parental transcriptome reference and indexed it with HISAT2 version 2.2.1[58]. We mapped paired-end RNA-seq reads to the combined transcriptome reference with HISAT2[58]. We disabled spliced alignment, soft clipping, mixed alignments, and discordant alignments, required a minimum alignment score of zero, and allowed HISAT2 to report up to 100 alignments per read, including secondary alignments. We converted the mapped SAM files to query-name-sorted BAM files using SAMtools version 1.9[59]. We counted only read pairs that mapped exactly and unambiguously to a single transcript. We retained alignments when reads were properly paired, both mates were mapped, the alignment was not supplementary, the edit distance was zero, and the CIGAR string contained no soft clipping, hard clipping, insertions, deletions, or skipped regions. We counted a read pair only when both mates had valid alignments to the same single transcript and no other transcript.

### Allele-specific expression (ASE)

We restricted ASE analyses to high-confidence one-to-one orthologs between CS and Paragon with transcript lengths differing by less than 100bp. We used parental RNA-seq samples to identify orthologs that could be assigned unambiguously to the expected parental haplotype. For each CS parent sample, we set missing counts to zero and retained genes with more than three reads assigned to the CS transcript and zero reads assigned to the Paragon transcript. We repeated the reciprocal filtering in Paragon parent samples, retaining genes with more than three reads assigned to the Paragon transcript and zero reads assigned to the CS transcript. We retained genes that passed these filters in all three CS parents and all three Paragon parents. This produced a set of one-to-one orthologs for which parental reads showed no detectable cross-cultivar haplotype mapping.

For each CS×P and P×CS hybrid, we retained genes that passed the parent-based filters described above. We first tested whether allelic read counts differed between the reciprocal hybrids. For this, we summed CS and Paragon allele counts to obtain total expression counts per hybrid sample and retained genes with CPM≥2 in at least four samples. We fitted a limma-voom model using TMM normalization factors estimated from total expression counts. The model included a sample effect, an allele effect, and an allele-by-cross-direction interaction. We identified genes with significant reciprocal-cross effects using the allele-by-cross-direction term at FDR<0.05 and removed these genes from subsequent ASE analyses. We then modeled allele-specific counts with a design including sample and allele effects, which compared CS and Paragon allele counts while accounting for paired counts from the same hybrid sample. We estimated TMM normalization factors from total expression counts and applied these factors to the allele-specific count matrix. We classified genes as showing significant ASE (H) when the allele effect had FDR<0.05 and |logFC| >1.

### Regulatory Divergence Classification

We tested the *trans*-regulatory contrast (T) by determining whether the parental expression ratio differed from the allelic expression ratio in the hybrids using two complementary approaches. In the first approach, we summed uniquely assigned transcript counts across parental replicates and allele-specific counts across hybrid replicates. We then used Fisher’s exact test to determine whether the CS:Paragon expression ratio differed between the parents and hybrids. Because this pooled-count approach does not account for variation among biological replicates[31], we also used a linear-model based analysis. For this, we summed the CS and Paragon allele-specific counts to obtain total counts for each hybrid library and calculated TMM normalization factors using the parental and total hybrid counts. We assigned each hybrid library’s size and normalization factor to both alleles. We then fitted a four-group linear model comprising CS parents, Paragon parents, CS alleles in hybrids, and Paragon alleles in hybrids. To account for the paired allele-specific measurements from each hybrid individual, we included hybrid-specific fixed effects, with one hybrid used as the reference level to maintain a full-rank design matrix. After voom transformation, we fitted gene-wise linear models using limma and tested the contrast:

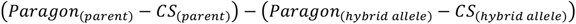

We applied empirical Bayes moderation to the gene-wise variances. For both approaches, we applied Benjamini–Hochberg correction and considered T significant at an adjusted *P*-value below 0.05.

Using the approach detailed in McManus *et al*.[30], we classified genes into regulatory divergence categories based on significant differences in parental expression (P), ASE (H), and *trans* effects (T). We assigned directionality using the fold changes. Genes were classified as *cis* only (P and H significant, T not significant), *trans* only (P and T significant, H not significant), *cis* + *trans* (all significant with concordant direction), *cis* × *trans* (all significant with opposing direction), compensatory (H and T significant but not P), or conserved (no significant effects). Genes that did not meet these criteria were classified as ambiguous.

### Homoeolog Expression Bias (HEB) Classification

We converted Ramírez-González *et al*.[14] triad annotations based on IWGSC RefSeq v1.1 to v2.1 using URGI Versailles correspondence tables (https://wheat-urgi.versailles.inra.fr/Seq-Repository/Assemblies) and retained 1:1:1 A:B:D homoeolog triads. For HEB analyses, we included triads with at least one expressed homoeolog (>0.5 transcripts per million in at least three samples) and required all three homoeologs to be detectable based on the criteria used for the differential gene expression analysis. We performed post hoc comparisons with Tukey’s honestly significant difference test in agricolae[60] following a one-way analysis of variance. To test whether triads with all three homoeologs differentially expressed could arise from sampling alone, we constructed a 6,591×3 triad matrix and randomly sampled 22.3% of cells (*i.e*., homoeologs) without replacement. We then counted triads in which all three homoeologs were sampled under this null model and repeated the procedure 10,000 times. We plotted the HEB distance of each hybrid observation from each parental centroid using ggpointdensity v0.2.1[61], with local point density estimated by two-dimensional kernel density estimation.

### Bisulfite Read Mapping and Methylation Calling

We trimmed paired-end bisulfite reads with Trimmomatic version 0.39[44] using the same parameters as above, but with a custom adaptor file. We indexed the IWGSC RefSeq v1.1 reference genome[28] and mapped reads with Bismark version 0.24[62] using HISAT2 and the following default parameters: --score-min L,0,-0.2, --ignore-quals, --no-mixed, --no-discordant, and --no-softclip. The availability of a whole genome comparative alignment between this reference and the Paragon GCA949126075v1 reference in Release 113 of Ensembl[57] allowed us to extract single nucleotide differences between the two cultivars. We used the URGI split version of the reference genome to enable memory-efficient alignment. We removed PCR duplicates and extracted cytosine-context methylation calls. We generated genome-wide cytosine reports with a minimum coverage of one read, retaining CG, CHG, and CHH calls. We merged replicate methylation counts for each genotype by summing methylated and unmethylated reads at matching sites, then separated calls into CG, CHG, and CHH contexts. For CG and CHG, we collapsed symmetric cytosine pairs across strands by summing the counts, using the plus-strand cytosine as the representative coordinate. Symmetric CG pairs were opposite-strand cytosines at consecutive positions, whereas symmetric CHG pairs were separated by two bases. We only kept sites that were typed in all three genotypes. We converted coordinates from the chromosome-split reference back to full chromosome coordinates.

We removed methylation sites overlapping single nucleotide differences between CS and Paragon, as they can confound bisulfite-based methylation calls in the hybrid. We used a custom script to extract single nucleotide differences between IWGSC RefSeq v1.1 and GCA949126075v1 from MAF alignments obtained from Ensembl Plants and generated BED outputs using v1.1 CS coordinates. For CG and CHG sites, we tested the corresponding two- and three-base contexts, respectively, for overlap with single nucleotide differences. We then used BEDTools version 2.31[63] to retain only CG and CHG methylation sites whose sequence contexts did not overlap any single nucleotide differences. Because of memory constraints, we used a custom Python script to remove CHH sites whose trinucleotide context overlapped single nucleotide differences. We defined the context as the cytosine and two downstream bases on the positive strand and the two upstream bases and cytosine on the negative strand.

### Annotation of Gene Body Methylation

We used a modified implementation of Muyle *et al*.[64] to identify gbM genes. For each genotype, we retained genes with at least 20 sites in each cytosine context, with each site covered by more than two reads. We classified individual sites as methylated using a one-sided binomial test with a non-conversion probability of 0.002, as reported to us by the sequencing provider, and a Benjamini–Hochberg-adjusted *P*-value below 0.001. For each genotype and cytosine context, we calculated the CDS-wide background methylation frequency as the number of retained CDS cytosines classified as methylated divided by the total number of retained CDS cytosines. For each gene, we used a one-sided binomial test to determine whether the number of methylated cytosines in each context exceeded that expected from the corresponding CDS-wide background frequency. We classified genes as gbM when they showed significant CG methylation enrichment at an Benjamini–Hochberg adjusted *P*-value of 0.05 or lower but no significant CHG or CHH enrichment. We classified genes as unmethylated (UM) when they showed no significant enrichment in CG, CHG, or CHH methylation. We excluded genes with other methylation patterns or without valid tests in all three contexts. We only retained only genes classified consistently as gbM or UM in CS, CS×P, and Paragon.

### Associations between Gene Body Methylation and Expression

We retained genes with transcripts per million (TPM) ≥1 in at least three out of nine libraries. The nine libraries consisted of three biological replicates each of CS, Paragon, and CS×P. We then calculated the mean TPM across the three replicates for each genotype. We used χ² tests to determine whether gbM frequency differed among the three subgenomes and whether methylation status (gbM or UM) was associated with expression class (additive, dominant, transgressive, or non-DE). We compared log₂(TPM+1) between gbM and UM genes using a Wilcoxon rank-sum test. We performed Tukey’s honestly significant difference test following a one-way analysis of variance to compare HEB between triads with varying numbers of gbM genes using agricolae[60].

## Availability of data and materials

RNA-seq and bisulfite-sequencing data are available in the NCBI Short Read Archive (https://www.ncbi.nlm.nih.gov/sra) under BioProject PRJNA1462369. Scripts and data are available at Zenodo (https://doi.org/10.5281/zenodo.21774265) and Figshare (https://doi.org/10.6084/m9.figshare.32144041).

## Competing interests

The authors declare that they have no competing interests.

## Funding

Funding from the Leonhard Lorenz-Stiftung and the European Commission’s Marie Skłodowska-Curie Actions (Grant ID: 101149103) supported the RNA-sequencing and bisulfite-sequencing, respectively.

## Acknowledgements

We thank the late Prof. Philippa Borrill for providing seeds and resources, and Emilie Knight for assistance with plant growth. This work was supported by the de.NBI Cloud within the German Network for Bioinformatics Infrastructure (de.NBI) and ELIXIR-DE (Forschungszentrum Jülich and W-de.NBI-001, W-de.NBI-004, W-de.NBI-008, W-de.NBI-010, W-de.NBI-013, W-de.NBI-014, W-de.NBI-016, W-de.NBI-022).

## Supplementary Information

Figures S1-S11

## Notes

### Competing Interest Statement

The authors have declared no competing interest.

### Summary of Updates

1) Removed the epigenetic inheritance results and refocused the manuscript on changes in gene expression and homoeolog expression bias following intraspecific hybridization, while retaining the gene body methylation results. 2) Applied a twofold-change threshold for identifying differential expression rather than 1.5-fold as before. 3) Removed all cytosine sites overlapping SNPs and adopted clearer filtering criteria and statistical methods for identifying gene body methylation. 4) Corrected an error in the trans test used to assign regulatory divergence categories. We now compare expression divergence between the parents with parental allelic expression divergence in the hybrids. We also added a separate linear-model-based trans test that is less sensitive to inter-individual variation. 5) Revised the title, abstract, introduction, and discussion to produce a more focused manuscript on gene expression changes in the intraspecific hybrid.

