## Supplementary Information for "Nonadditive gene expression and reduced homoeolog expression bias in an intraspecific hexaploid wheat hybrid"

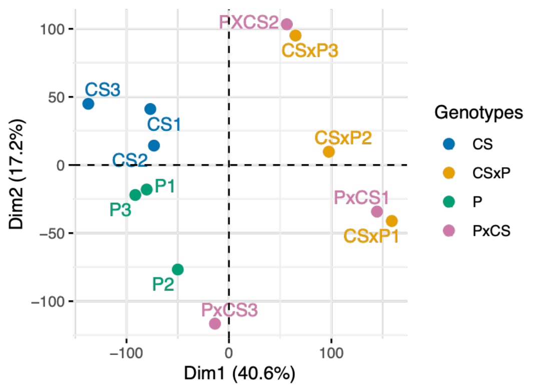


**Figure S1**. Principal component analysis (PCA) using read counts for Chinese Spring (CS) and Paragon (P) parental lines and six hybrid genotypes from reciprocal crosses.

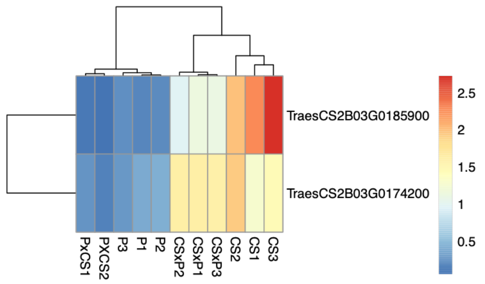


**Figure S2**. Counts per million (CPM) for genes where cross direction had a significant effect on expression (limma–voom empirical Bayes moderated t-test with FDR<0.05). CPM values were normalized by dividing each gene's expression by its mean across all samples.

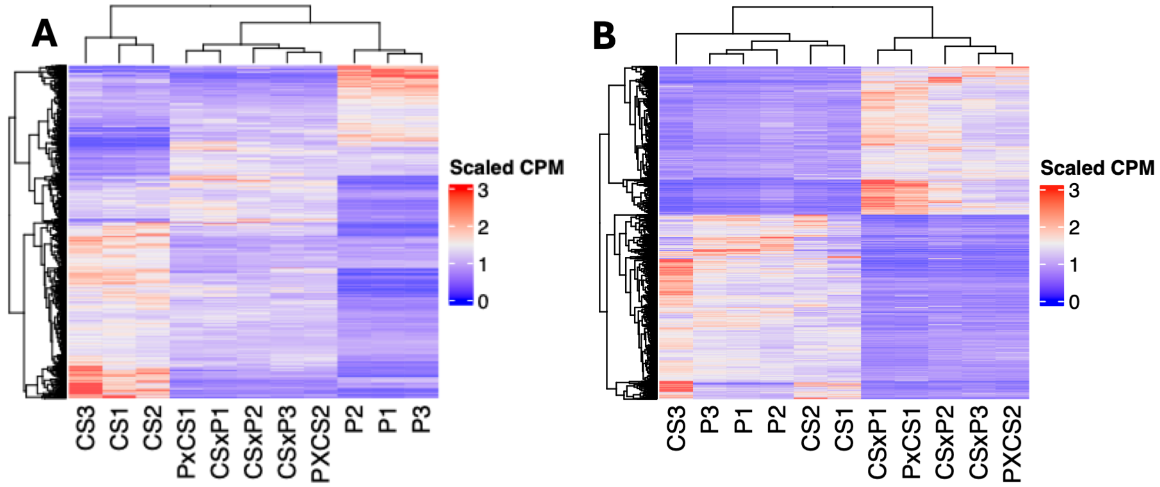


**Figure S3**. Counts per million (CPM) gene expression estimates for genes that were significantly differentially expressed **(A)** between the Chinese Spring (CS) and Paragon parents and **(B)** between mid-parental estimates and the hybrids. Differential expression was identified using a limma–voom empirical Bayes moderated t-test (≥2-fold expression difference and FDR<0.05). CPM values were normalized by dividing each gene's expression by its mean across all samples.

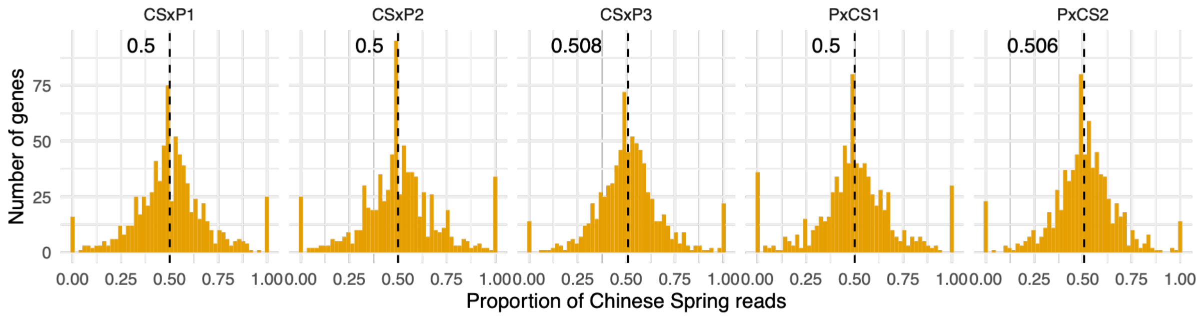


**Figure S4**. Proportion of reads mapping exactly and uniquely to the Chinese Spring (CS) haplotype in CS–Paragon hybrids. Analyses were restricted to one-to-one parental orthologs for which parental RNA-seq reads showed no cross-cultivar haplotype mapping.

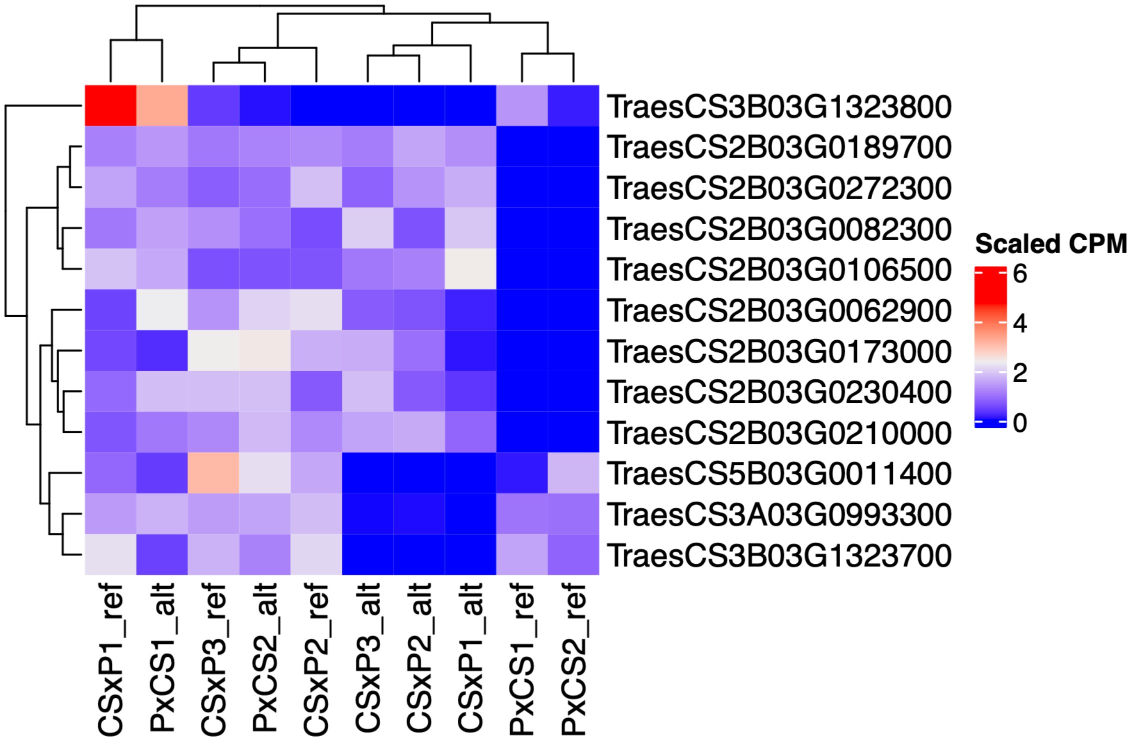


**Figure S5.** Counts per million (CPM) gene expression estimates for genes showing differential allele-specific expression between reciprocal hybrid crosses between Chinese Spring (CS) and Paragon. Differential expression was identified using a limma–voom empirical Bayes moderated t-test (FDR<0.05). CPM values were normalized by dividing each gene's expression by its mean across all samples.

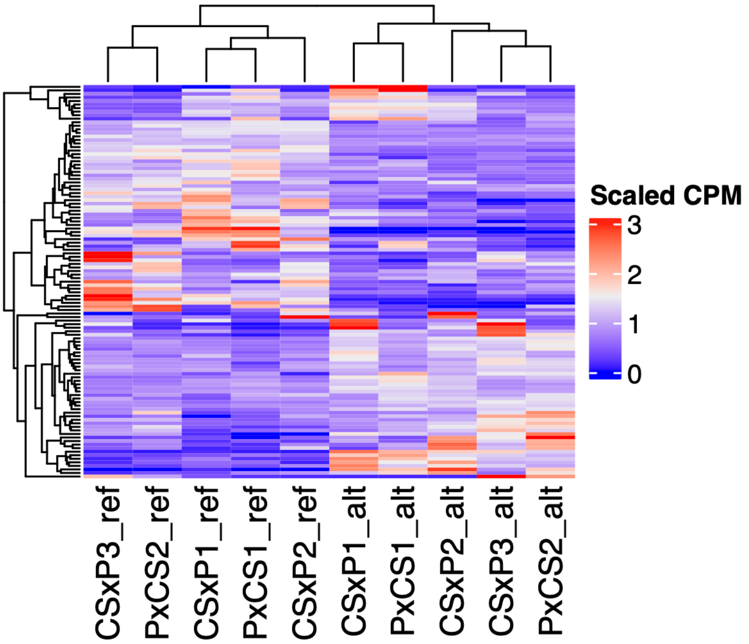


**Figure S6**. Counts per million (CPM) gene expression estimates for genes that showed significant allele specific expression in hybrids between Chinese Spring (CS) and Paragon. Differential expression was identified using a limma–voom empirical Bayes moderated t-test (≥2-fold expression difference and FDR<0.05). CPM values were normalized by dividing each gene's expression by its mean across all samples.

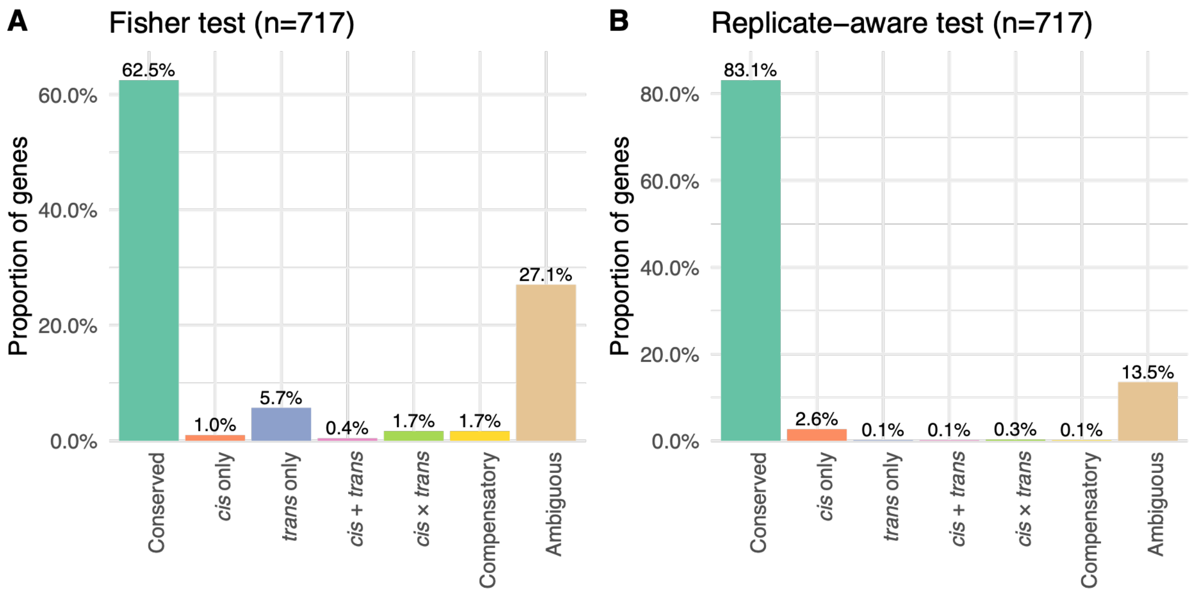


**Figure S7**. Regulatory category assignments for genes. **(A)** Classification using the binomial and Fisher's exact test. **(B)** Classification using the linear model method. n indicates the total number of classified genes.

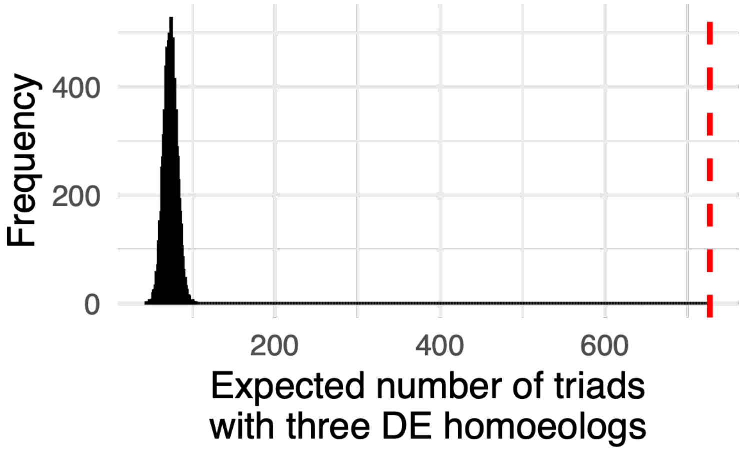


**Figure S8**. Expected number of triads in which all three homoeologs are differentially expressed by chance alone, if 22.3% of genes are differentially expressed. The histogram shows results from 10,000 replicate simulations. The red line indicates the observed number of triads in which all three homoeologs are differentially expressed in hybrids relative to mid-parental estimates.

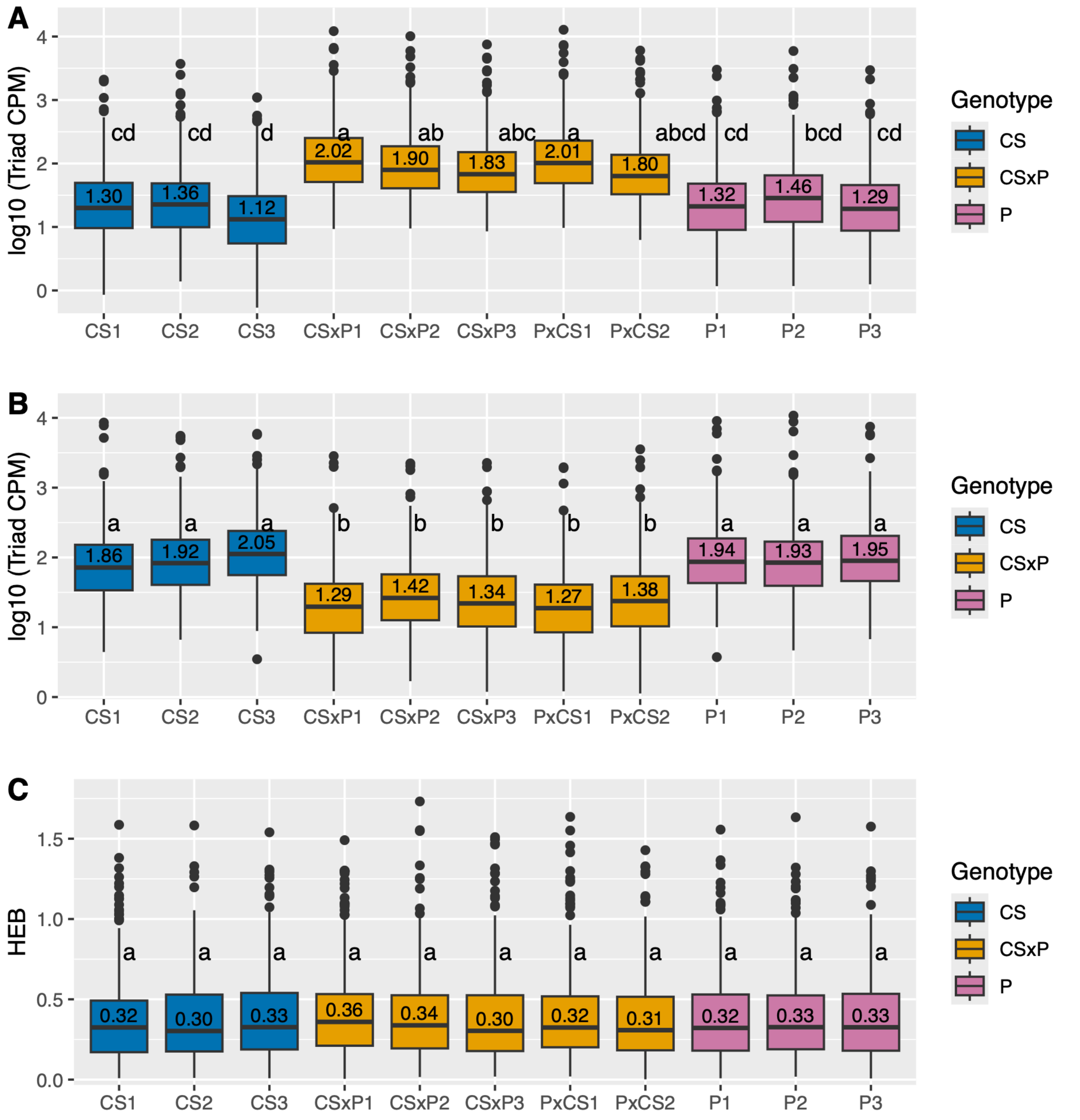


**Figure S9**. Triad expression levels and homoeolog expression bias (HEB; measured as the coefficient of variation) for triads in which all three homoeologs are differentially expressed in hybrids relative to mid-parental estimates for Chinese Spring (CS), Paragon (P), and CS×P hybrid genotypes. **(A)** Counts per million (CPM) of triads overexpressed in hybrids. **(B)** CPM and **(C)** CV of triads underexpressed in hybrids. Tukey's HSD tests were performed following one-way ANOVA of triad CPM or CV. Samples sharing the same letter are not significantly different (*P*>0.05).

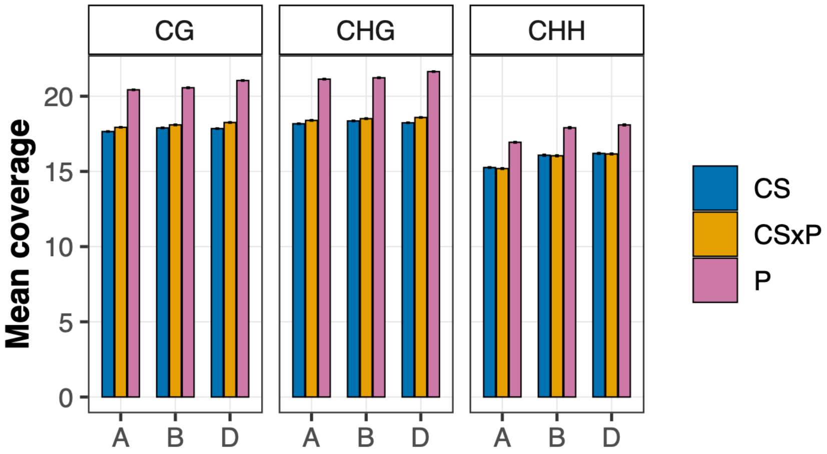


**Figure S10.** Average number of reads per site (coverage) for CG, CHG, and CHH contexts for the A, B and D subgenomes for Chinese Spring (CS), Paragon (P), and CS×P hybrid genotypes. Error bars indicate confidence intervals.

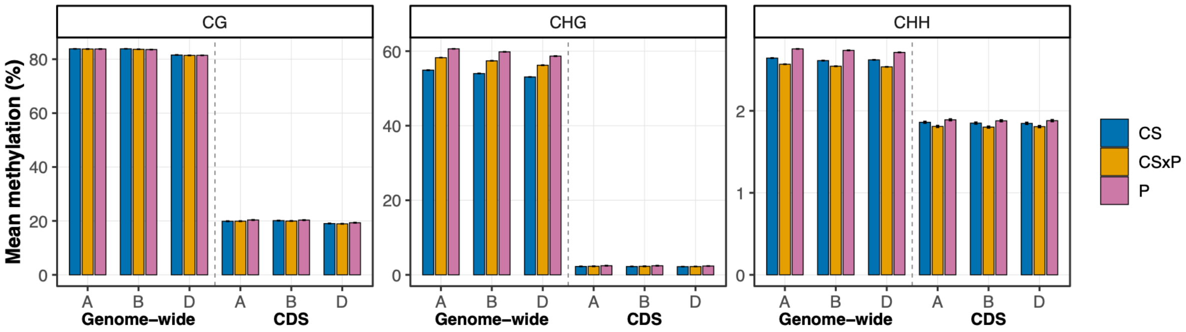


**Figure S11.** Mean CG, CHG, and CHH methylation levels genome-wide and in coding sequence (CDS) intervals for Chinese Spring (CS), Paragon (P), and CS×P hybrid genotypes. Error bars indicate confidence intervals.
